# Whole genome sequencing with AVITI and NovaSeq X Plus reveals comparable performance with contextual biases

**DOI:** 10.1101/2025.10.10.681584

**Authors:** Pontus Höjer, Johannes Alneberg, Pär Lundin, Tom Martin, Julia Hauenstein, Helena Fällmar, Magnus Lindell, Christian Natanaelsson, Susana Häggqvist, Adam Ameur, Jessica Nordlund, Robert Månsson Welinder

## Abstract

Element Biosciences’ avidity sequencing has emerged as a competing technology to Illumina’s short read sequencing platform. Prior benchmarks of avidity sequencing have not included the latest Illumina NovaSeq X/X Plus instruments with XLEAP chemistry. Here, we have run PCR-free whole genome sequencing on four human tumor cell lines using both Illumina NovaSeq X Plus and Element AVITI instruments. AVITI showed low duplication rates and reported higher base qualities, the latter contributed to improved mapping confidence and fewer spurious variant candidates. Both platforms were found to be highly comparable when benchmarking variant calling, with AVITI only providing a minor improvement on INDELs at lower coverages. Stratifying by genomic context revealed further differences, where AVITI genome coverage and variant calls were superior in high GC regions while being inferior in GC homopolymers. Error rate analysis highlighted further differences between the platforms, in particular AVITI in some instances displayed an increased error rate on read 2 related to short fragments. AVITI error rate was also found to be more stable downstream of repetitive regions, except for GC homopolymers. We further found that AVITI sequencing was sensitive to G-quadruplex motifs. Overall, despite these identified differences, both platforms performed highly comparable for variant analysis.

## INTRODUCTION

Massively parallel sequencing has become a true workhorse for genomics research since its introduction in the early 21st century. The ability to read large numbers of DNA sequences has many applications, most prominently in medicine and biology. While initially multiple sequencing technologies competed in this space (1), Illumina’s (Solexa’s) ‘sequencing by synthesis’ (2) rapidly became the dominant technology (3). Later years have however seen increased diversification coupled to the emergence of new competing short read (<1000 bases) platforms (4) such as Element Biosciences’ AVITI system that relies on novel avidity sequencing technology (5).

Avidity sequencing, while similar to Illumina in many regards, comes with several important differences. Both sequencing technologies amplify a single DNA template into ensembles of localized copies on a flow cell surface. These ensembles are then read by incorporating and imaging nucleotide bases one by one along the template, starting from a common primer sequence. For amplification, Illumina relies on PCR-like bridge amplification to generate clusters of clonal sequences (2). Element instead relies on rolling circle amplification (RCA) to generate long strands containing multiple template copies that are collapsed into a structure referred to as a polony (5). Unlike PCR, RCA avoids propagation of errors since it copies strictly from the original template (6). For base incorporation and reporting, Illumina uses nucleotides with a fluorescently-labelled reversible terminator that are incorporated and imaged before removing the terminator to allow the next base to be incorporated and read (7). In avidity sequencing, reporting is performed by avidites that are bound, imaged, and subsequently removed to allow incorporation of an unlabeled reversibly-terminated nucleotide. Avidites consist of multiple nucleotides of a common type, flexibly chained to a fluorescent core. This setup allows multiple nucleotides from the same avidite to bind complementary bases within the same polony. Binding of the avidites to multiple bases increases the signal retention to improve signal-to-noise ratios (5). This also limits the impact of phasing, a common issue in ensemble-based sequencing, where some template strands lag or advance out of sync (8). Furthermore, bulky fluorescent-labels, as used by Illumina, can exacerbate phasing by impeding base incorporation (9).

Element sequencing has previously been benchmarked against Illumina for the purpose of whole genome sequencing (WGS) (5, 10), but to our knowledge no independent comparison of the technologies has been made. Furthermore, published comparisons were not made to include the latest large-scale sequencing instruments released by Illumina (NovaSeq X/X Plus) using updated X-LEAP chemistry, reported to yield higher base qualities (11). For this purpose, we performed WGS on four human cell lines, sequencing the same PCR-free libraries on both Illumina NovaSeqX Plus (NovaSeqX+) and Element AVITI instruments. Using these datasets, we evaluate multiple characteristics of the technologies for WGS, including variant calling performance and sequencing error modes in various contexts. Altogether, we find the technologies highly comparable, but with specific differences related to among other things genomic context and error modes. Specifically analyzing the rare disparities, we show that Element sequencing is affected by G-quadruplex motifs.

## METHODS

### PCR-free library preparation and sequencing

Multiple myeloma (MM) cell lines MM1.S (MM1S), OPM2 and KMS12BM were cultured in RPMI-1640 medium with HEPES and GlutaMAX (Thermo Fisher Scientific, #72400021), supplemented with 10% FCS (Sigma, #F7524) and 100 U/ml Penicillin + 100μg/ml Streptomycin solution (Cytiva, #SV30010). Genomic DNA was extracted using Monarch® HMW DNA Extraction Kit for Cells & Blood (NEB, #T3050) following the manufacturer’s instructions. The acute lymphoblastic leukemia (ALL) cell line REH was cultured in RPMI-1640 medium (Sigma, #R0883) supplemented with 2 mM L-Glutamine (Sigma, #G7513), 100 U/mL Penicillin and 100μg/mL Streptomycin (Sigma, #P0781), and 10% heat-inactivated fetal bovine serum (HIFBS) (Sigma, #F9665) as previously described(12). Genomic DNA from REH cells was extracted using the Nanobind CBB Big DNA Kit (Circulomics, NB-900-001-01) following the instructions in “Nanobind UHMW DNA Extraction – Cultured Cells Protocol” (Circulomics, EXT-CLU-001). Library preparation was performed using the TruSeq DNA PCR-Free kit (Illumina, #20015962) according to the manufacturer’s instructions. For each cell line, three replicate libraries were generated, each with unique UDI indexes, resulting in a pool of 12 libraries.

The library pool was sequenced on Illumina NovaSeq X Plus and Element AVITI instruments. Illumina NovaSeq Plus sequencing (performed at the Swedish National Genomics Infrastructure, NGI) was done using two lanes using a NovaSeq X Series 10B 300-cycle Reagent Kit (Illumina, #20085594) with a 2x151 bp read setup. The lanes were loaded with 140 pM and 160 pM of denatured library, respectively. The run utilized control software version 1.2.0.28691, and data conversion was done using bcl2fastq version 2.20.0.422. Element AVITI sequencing was initially done using Element Adept Rapid PCR-Free Kit for library conversion, followed by sequencing (performed as a service by Element Biosciences, USA, San Diego) on two Cloudbreak (CB) High Output flow cells (2x150 bp, #860-00003). The run was performed using an AVITI instrument and demultiplexing with bases2fastq v1.6.1.1089765930. Element AVITI sequencing was also done (performed by NGI) using one Cloudbreak Freestyle (CB FS) High Output flow cell (2x150 bp, #860-00013) to load the pool without the need for library conversion. Each lane was loaded at 10 pM and 12 pM respectively. The run was performed using an AVITI instrument running AVITI OS v2.6.2 and demultiplexing with bases2fastq v1.8.0.1260801529.

### Sequencing data processing

Primary analysis was performed using the nf-core/sarek pipeline (v2.4.3) (13, 14) built on Nextflow (v24.4.2) (15), mapping with BWA MEM (v0.7.17.post1188) (16) to the GRCh38 human reference genome (https://ftp-trace.ncbi.nlm.nih.gov/giab/ftp/release/references/GRCh38/GRCh38_GIABv3_no_alt_analysis_set_maskedGRC_decoys_MAP2K3_KMT2C_KCNJ18.fasta.gz) and marking duplicates with picard MarkDuplicates (gatk v4.5.0.0, picard v3.1.1) (17). The pipeline was configured to skip the default fastp (v0.23.4) (18) read trimming, except for capping read length to the same 150 bp across the datasets. Base recalibration was further disabled (option “--skip-tool baserecalibrator”) to preserve the original base qualities.

Information about insert size, base qualities, and mapping quality (MAPQ) was gathered from samtools stats (v1.19.2) (19) that is part of the nf-core/sarek pipeline.

### Duplicate reads investigation

Reads marked as duplicates by gatk/picard MarkDuplicates were investigated to see if the origin could be optical or from complementary strands. For this, duplicates were re-marked with MarkDuplicates but with the option “--TAG_DUPLICATE_SET_MEMBERS true” to label reads involved in a duplicate set. These labeled reads were then extracted and each set investigated using a custom script (<u>duplicates.py</u>), running Python (v3.12.7) with the pysam (v0.22.1) package. For each combination of read pairs in a duplicate set, the distance between the pairs and the pair’s relative order was computed. Pairs within 100 distance units were classified as “proximal”. Pairs where the reads map in opposite directions were classified as “complementary”. Each combination of classifications were tallied.

### Differential coverage

Datasets were first downsampled to 10X coverage using samtools view (v1.19.2) with option “-s {fraction}”. The fractions were based on the mean coverage acquired using mosdepth (v0.3.8) (20). We generated d4 format files (21) using mosdepth (v0.3.10) for reads with a MAPQ of at least 60 to look up the coverage per base across multiple genomic regions. The d4 file format allows for fast coverage queries across multiple intervals. To assay coverage across different genomic contexts, we used the Genome in a Bottle (GIAB) GRCh38 genome stratifications (v3.5, https://ftp-trace.ncbi.nlm.nih.gov/giab/ftp/release/genome-stratifications/v3.5/) (22) with d4tools (v0.3.10 with d4 library v0.3.9) to get coverage distributions for each stratification. The mean coverage for each stratification and sample was calculated from the distribution. Stratifications covered by less than 10, 000 bases total were excluded from further analysis. We further excluded stratifications in the subcategories “Ancestry”, “FunctionalTechnicallyDifficult”, and “GenomeSpecific”, as well as relating to the X and Y chromosomes. The top 15 most variable stratifications were selected based on their variance in mean coverage across the AVITI and NovaSeqX+ runs for each cell line. Finally, the relative stratification coverage difference to the overall autosomal coverage (stratification “GRCh38_AllAutosomes.bed.gz”) was calculated.

### Variant calling and benchmarking

Element and Illumina variants were called with DeepVariant (v1.5.0) (23) on chr20 using the model “WGS”. Read were downsampled to 10X, 15X, 20X, 25X, 30X, 40X, 50X using “--make_examples_extra_args downsample_fraction={fraction}” using fractions calculated from the mean chr20 coverage acquired from mosdepth (v0.3.8).

Benchmarking on chr20 was performed using hap.py (v0.3.8-17-gf15de4a) (24) with the RTG-tools vcfeval engine with option “--engine=vcfeval”. PacBio DeepVariant PASS calls were used as truth set, and with option “--preprocess-truth” enabled. Benchmarking was limited to high-confidence regions (see section below for how these were generated). To query different genomic contexts, GIAB stratifications (v3.5) were used. Variant calling performance was assayed using the F1-score metric calculated as below.

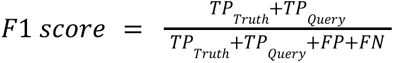

Here, true positives (𝑇𝑃) are disambiguated into 𝑇𝑃_𝑇𝑟𝑢𝑡ℎ_, the number of variants in the truth-set that match the query, and 𝑇𝑃_𝑄𝑢𝑒𝑟𝑦_, the number of variants in the query that match the truth-set. False positives (𝐹𝑃) are query variants not matched in the truth-set. False negatives (𝐹𝑁) are truth-set variants that are missing from the query.

### PacBio sequencing and variant calling

PacBio HiFi sequencing libraries for MM1S, OPM2, and KMS12BM were prepared using the same high molecular weight genomic DNA source as the short read libraries and the SMRTbell prep kit 3.0 (Pacific Biosciences, #102-141-700). Sequencing was done on the PacBio Revio system. The PacBio reads were mapped to GRCh38 using pbmm2 (v1.10.0, https://github.com/PacificBiosciences/pbmm2). Sequence length and base quality was assessed using sequali (v1.0.2) (25). Genome coverage was computed using mosdepth (v0.3.8). Small variants were called using DeepVariant (v1.5) using the “PACBIO” model. Structural variants were called using PBSV (v2.9.0).

### Generation of high-confidence benchmarking regions

We generated high-confidence benchmarking regions for each MM cell line. First, across all datasets, we removed regions with low/high coverage. For this purpose, mosdepth (v0.3.8) was used to query the coverage in 100 bp windows, limiting to reads with at least MAPQ 60 for Illumina/Element and MAPQ 20 for PacBio. Any windows with less than 10X coverage or more than two times the median dataset coverage were excluded. The remaining Illumina/Element regions were merged using bedtools (v2.30.0) (26) for each cell line.

Complex variant regions were excluded by finding regions with two or more variants within 10 bp in the filtered PacBio DeepVariant calls and extending these by 50 bp on each side. We also excluded regions within 50 bp of SVs greater than 50 bp called with PBSV.

Inspired by previous benchmarking efforts by the GIAB consortium (27, 28) we further excluded contigs shorter than 500kb (GRCh38_contigs_lt500kb) and regions close to reference gaps (GRCh38_gaps_slop15kb), relying on GIAB stratifications v3.5. We further excluded regions with long homopolymers (GRCh38_SimpleRepeat_imperfecthomopolge11_slop5 and GRCh38_SimpleRepeat_homopolymer_ge12_slop5) that are problematic for PacBio HiFi reads (29, 30). Finally, we further removed any partially covered tandem repeats or homopolymers (defined by GRCh38_AllTandemRepeatsandHomopolymers_slop5) similar to Wanger et al. (28). Bedtools (v2.30.0) (26) and bcftools (v1.18) (31) were used extensively for generating the high-confidence regions.

### Publicly available avidity sequencing datasets

For the error rate evaluation, we in addition to our own data utilized publicly available Element avidity sequencing data, generated using different PCR-free library preparation kits and AVITI flowcells (Supplementary Table 1). For five datasets, we downloaded BAM files with GRCh38 mapped reads. The HG002 Cloudbreak Freestyle dataset that was only available as FASTQ files was downloaded and mapped to GRCh38 using the nf-core/sarek pipeline (v2.4.3) and the same configurations as described above.

### Error rate investigation

Reference-based error rate per cycle was evaluated using samtools (v1.19.2) for read 1 and read 2 separately on chr20. Only reads with a mapping quality of at least 60 were included. Reads were selected using samtools view and statistics were calculated using samtools stats. Bases and substitutions versus the reference were aggregated for each cycle to calculate the error rate. The error rates were smoothed using a 5 bp rolling average across cycles.

We evaluated the overlap-based error rate per cycle using fraguracy (v0.2.4, https://github.com/brentp/fraguracy) with the option “--bin-size 1”. The error rates were smoothed using a 5 bp rolling average across cycles. To calculate the median error rate per read, only bases at positions 50 to 60% into the read were considered, similar to Stoler & Nekrutenko (32).

For the insert-size based error evaluation, reads were selected by their template length in bins of 50 bp using samtools view for read 1 and 2 separately. Bases and errors versus the reference were calculated using samtools stats as above.

To evaluate error rate by genomic context, we relied on the “stack_reads_by_interval.py” script from Arslan et al. (commit 701be39, https://github.com/Elembio/AvidityManuscript2023) (5). Repetitive regions in GRCh38 we acquired from GIAB (v3.5). A control set of one million 30 bp random regions was generated using bedtools random with options “-l 30 -n 1000000”. Briefly, reads on chr20 with MAPQ >=60 overlapping the regions were extracted using samtools. These reads were then piled up across each region by strand to identify substitutions versus the reference using the “stack_reads_by_interval.py” script. The error rate for the bases upstream, inside and downstream of the regions was computed. For the upstream and downstream error rate, only the 50 bases closest to the region were used.

### G-quadruplex investigation

The G-quadruplex (G4) investigation relied on predicted G4 (pG4) motifs generated using pqsfinder (v2.0.1) (33) with default options on GRCh38 downloaded from https://pqsfinder.fi.muni.cz/genomes.

To study the overlap between extensively soft-clipped alignments and pG4 motifs, we extracted reads with at least 10 soft-clipped bases using samtools (v1.19.2). Only autosomal reads with a MAPQ greater than or equal to 60 and a template length (insert size) greater than 150 bp were included. These reads were converted to BED format using bedtools (v2.30.0) bamtobed and split by strand information (forward or reverse). For each strand and sample, the read spans were extended by 10 bp on the ends using bedtools slop and then merged using bedtools merge, counting the number of overlapping spans. Only spans with at least two soft-clipped reads were kept. For each strand and cell line, only spans shared across the AVITI CB and AVITI CB FS runs were kept. These spans were further filtered to only keep those that were shared across at least two cell lines. This yielded locations of prominent soft-clipping across the AVITI datasets on each strand. Soft-clipping is common for reads that overlap structural variations. To exclude such sites, locations with overlapping soft-clipped reads on both strands were removed. As an additional layer of filtration, common soft-clipped sites in the NovaSeq X Plus datasets were subtracted from the AVITI sites to get sites specific to the AVITI data. Overlap between the AVITI soft-clipped sites and pG4 sites by strand was queried using bedtools intersect.

We further looked at base quality scores and mismatched bases for reads overlapping pG4 sites. G4 sites might be proximally spaced, leading to a read spanning multiple sites. Therefore, we focused on isolated pG4 sites, removing any sites within 300 bp of each other using bedtools (v2.30.0). Any pG4 sites longer than 50 bp were also excluded to only include sites that can be covered by a 150 bp read. Reads on chr20 overlapping the isolated pG4 sites were extracted using samtools (v1.19.2). Only reads that covered the pG4 sites by at least 5 bp on each side were included. Both the isolated pG4 sites and the overlapping reads were split by strand. Reads were piled up for each site expanded by 150 bp using samtools mpileup with options ‘--no-BAQ --min-BQ 0 -a -x’. From the pileups, base qualities and mismatched bases were collected for each position relative to the pG4 site on both strands using a custom Python script.

The error rate following G4 motifs was assayed using the stack_reads_by_interval.py script from Arslan et al. (commit 701be39, https://github.com/Elembio/AvidityManuscript2023) (5), similar to the error rate investigation detailed above.

### Data visualizations

Data visualizations were generated using Python (v3.12.7) with extensive use of packages jupyter (v5.7.2) (34), seaborn (v0.13.2) (35), pandas (v2.2.3) (36, 37), matplotlib (v3.9.2) (38), numpy (v2.1.3) (39) and scipy (v1.14.1) (40). Figures and illustrations were composed and visually adjusted for clarity using Affinity Designer (v2.6.0).

## RESULTS

To directly compare Element to Illumina sequencing, the same pool of PCR-free gDNA libraries was sequenced on both the Element AVITI and the Illumina NovaSeq X Plus sequencers (Figure 1a). Illumina-style PCR-free WGS libraries were prepared using DNA from four cell lines, three of multiple myeloma origin (MM1S, OPM2, KMS12BM) and one with acute lymphocytic leukemia origin (REH). For Illumina sequencing, the pool was loaded on two 10B flowcell lanes on a NovaSeq X Plus instrument. Two rounds of Element AVITI sequencing were performed. In the first round (referred to as AVITI CB), libraries were converted to circular Element-style libraries and loaded onto two CB flow-cells for sequencing. In the second round (referred to as AVITI CB FS), libraries were directly loaded on a single Element CB FS flowcell for on-board circularization and sequencing. A schematic overview of the different workflows can be seen in Supplementary Figure 1.

**Figure 1:**
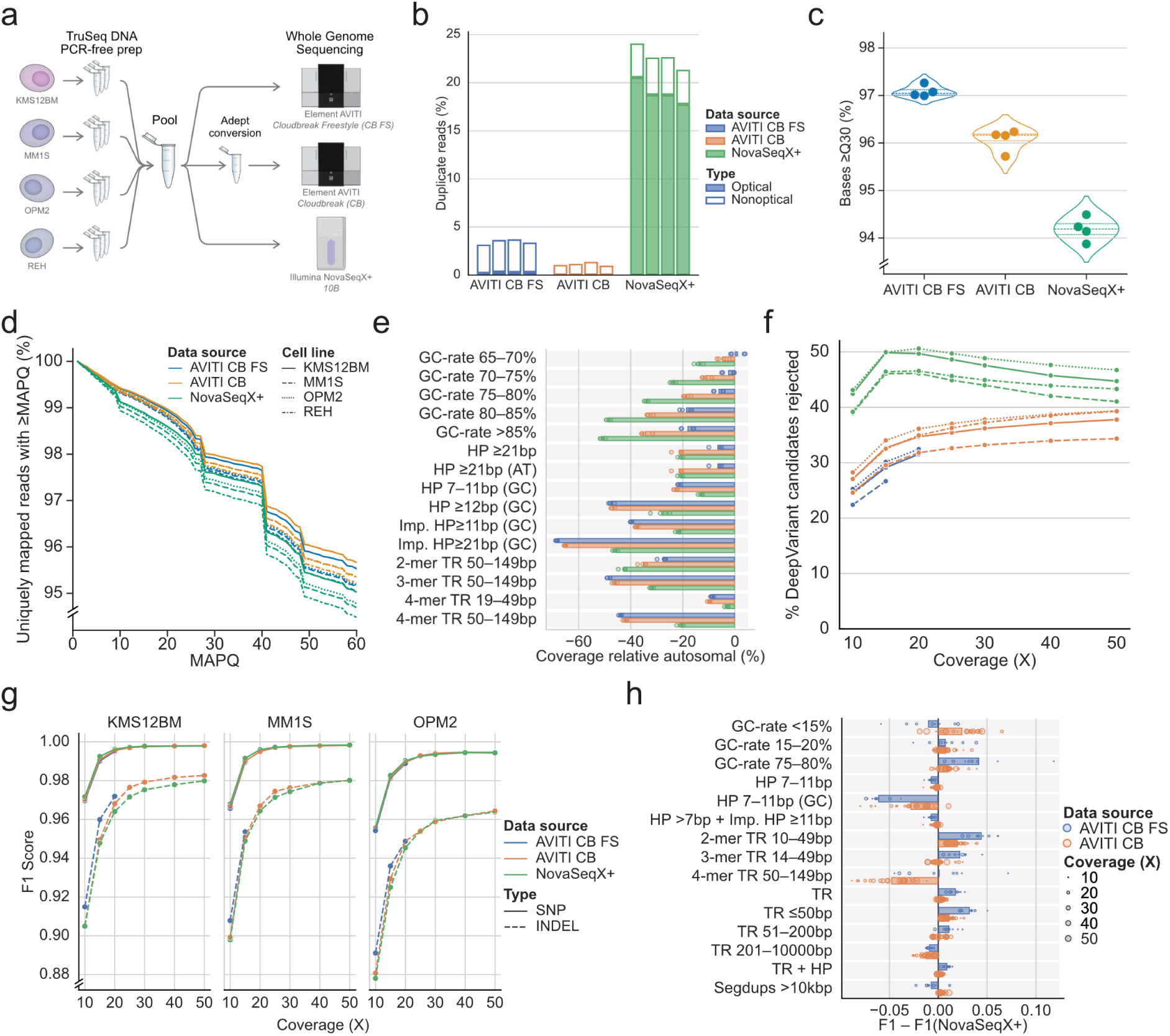
Comparative performance of Element and Illumina NovaSeq X Plus for WGS. (**a**) Study design. (**b**) Duplication rates by category. For the Illumina NovaSeq X Plus results it should be noted that one lane had an overall inflated duplication rate due to suboptimal loading, see Supplementary Table 2. (**c**) Reported base quality greater than or equal to Q30. (**d**) Mapping confidence for BWA-MEM on GRCh38 as measured by the percentage of all uniquely mapped reads with a particular mapping quality (MAPQ) or greater. (**e**) Differential coverage across the top 15 most variable genome stratifications. Difference measure relative to mean autosomal (“AllAutosomes”) coverage for the dataset. (**f**) Candidate small variant calls filtered out by DeepVariant across coverage from 10 to 50X. (**g**) F1 score for variant calling on chr20 for SNPs and INDELs at different levels of genome coverage, as reported by hap.py. PacBio HiFi DeepVariant calls were used as the truth set. Cell line REH was omitted as HiFi coverage was insufficient. (**h**) Relative F1 score for SNVs and INDELs combined comparing AVITI and NovaSeq X Plus across GIAB stratifications (v3.5) for the top 15 most variable stratifications. TR = tandem repeat, HP = homopolymer, Imp. HP = imperfect HP, N-mer = TR composed of N bp repeated element, SegDups = Segmental duplications.

Initial WGS analysis and quality control were performed using the nf-core/sarek pipeline (v2.4.3) and mapping to GRCh38 was done using BWA-MEM. Both read trimming and base quality recalibration were disabled. Mapping rates were high (>99%) across all samples (Table 1). Insert sizes were also similar across all datasets (∼400 bp), indicating no apparent size-dependent bias in fragments sequenced by either instrument (Supplementary Figure 2). Mean coverage was expectedly lower for the AVITI CB FS datasets (15.2 to 20.0X), while the other datasets had more than 40X coverage (Table 1).

**Table 1:** AVITI and NovaSeq X Plus statistics for cell line libraries and sequencing setups. CB = Cloudbreak, FS = Freestyle.

| Data Source | Cell line | Read Pairs (M) | Mean insert size (bp) | Mapped (%) | MAPQ=0 (%) | Mean Coverage (X) |
| --- | --- | --- | --- | --- | --- | --- |
| AVITI CB FS | KMS12BM | 190 | 392 | 99.9 | 4.0 | 17.7 |
| AVITI CB FS | MM1S | 191 | 397 | 99.9 | 4.2 | 17.7 |
| AVITI CB FS | OPM2 | 216 | 387 | 99.9 | 4.1 | 20.0 |
| AVITI CB FS | REH | 163 | 410 | 99.8 | 4.1 | 15.2 |
| AVITI CB | KMS12BM | 539 | 406 | 99.9 | 3.7 | 51.4 |
| AVITI CB | MM1S | 525 | 411 | 99.9 | 3.9 | 49.9 |
| AVITI CB | OPM2 | 526 | 400 | 99.9 | 3.9 | 50.1 |
| AVITI CB | REH | 475 | 425 | 99.9 | 3.9 | 45.3 |
| NovaSeqX+ | KMS12BM | 694 | 406 | 99.7 | 4.6 | 52.4 |
| NovaSeqX+ | MM1S | 687 | 412 | 99.7 | 4.9 | 51.0 |
| NovaSeqX+ | OPM2 | 787 | 400 | 99.7 | 4.8 | 58.4 |
| NovaSeqX+ | REH | 641 | 425 | 99.7 | 4.9 | 46.7 |

Increased duplication rates have been reported for other Illumina instruments compared to Element Avidity sequencing (5, 41). Unlike Element, Illumina clustering generates multiple copies of the original fragment that can seed adjacent wells in patterned flow-cells, generating optical duplicates. Indeed, NovaSeq X Plus duplication rates were found to be higher compared to the AVITI runs and primarily driven by optical duplicates (Figure 1b). In our results, the NovaSeq X Plus duplicate rates were partially inflated by nonoptimal loading in the lane with the lower loading concentration (140 pm) (Supplementary Table 2). However, even in the best NovaSeq X Plus lane, duplicates contribute to a substantial portion of reads (12-13%) (Supplementary Table 2). Element AVITI duplicates (<5% in all lanes) were primarily non-optical and seemingly originated from sequencing of the complementary strands of the original genomic DNA fragment (Supplementary Figure 3). These nonoptical duplicates were also slightly less frequent in AVITI CB than AVITI CB FS, possibly due to the CB library conversion that denatures and dilutes complementary strands before loading. Duplicate marking may thus be inadvisable for AVITI sequencing of PCR-free libraries.

Sequencing instruments generally report a phred quality score (Q) for each called base that estimates the probability of the base being called correctly. Avidity sequencing has been reported to give highly accurate base calls, with 85.4% of bases being >Q40 (less than one error per 10, 000 bases) (5). Similar values were found in the AVITI CB datasets reported here, with 84.0% to 85.4% of base qualities >Q40. The AVITI CB FS base qualities were even higher, with 87.7% to 88.5% of bases >Q40. Direct comparison to Illumina NovaSeq X Plus data is made complicated by the instrument reporting Q scores in four bins (bin 2, Q0-2; bin 12, Q3-17, bin 24, Q18-29; bin 40, Q30+; with Control Software v1.2.0), where base qualities ≥Q30 are reported in bin 40 (Supplementary Figure 3). We therefore relied on the fraction of bases ≥Q30 for comparisons against the AVITI datasets. The AVITI datasets had 95.7 to 97.3% of bases ≥Q30, compared to ∼94% for NovaSeq X Plus data (Figure 1c). In the AVITI data, the Q-scores were also slightly higher in the AVITI CB FS datasets, which may be due to the lower polony density (∼800 million per lane) compared to the AVITI CB runs (∼1000 million per lane).

### Comparison of mapping confidence, coverage biases and variant calling

To evaluate the effects of the higher accuracy base calls in the AVITI data, we investigated alignment to the GRCh38 human reference genome. Sequencing errors complicate genome mapping as they increase the number of plausible locations within the reference. Most mappers, including BWA-MEM, report a phred-scaled mapping quality (MAPQ) that measures the certainty of the read placement. Ambiguously mapped reads are given a MAPQ of zero. This was more commonly observed for NovaSeq X Plus reads (4.6 to 4.9% of mapped reads) compared to AVITI (3.7 to 4.2% of mapped reads). For uniquely mapped reads, the AVITI datasets generated higher MAPQ scores with BWA-MEM relative to NovaSeq X Plus (Figure 1d). These findings indicate that the increased base call accuracy of AVITI data contributes to improved read mapping confidence.

Looking further into the alignments, we investigated coverage biases by genomic contexts. Genomic contexts for GRCh38 were acquired from Genome in a Bottle (GIAB) stratifications (22). To measure the coverage bias, the difference in mean coverage per stratification to the autosomal coverage was calculated for each sample. Looking at the top variable stratifications, most coverage variation was observed in repetitive and high GC regions (Figure 1e). Compared to NovaSeq X Plus, the AVITI samples had higher coverage in high GC regions and 2-mer tandem repeats (TR) between 50-149 bp, while coverage was lower in 3- and 4-mer TRs as well as GC homopolymers (HPs) and GC imperfect HPs. AVITI CB and CB FS displayed mostly similar coverage, but AVITI CB FS had higher coverage in long (≥21bp) non-GC homopolymers (HPs) and in high GC regions.

Besides mapping, base quality also influences variant calling. For this, we used DeepVariant (23) v1.5, which includes a model trained on both Illumina and Element sequencing data. DeepVariant generates candidate variants, which the model then evaluates to either reject or keep the call. Candidate generation is highly permissible to increase sensitivity, requiring support of two reads as well as allele frequencies of at least 6% (SNPs) or 12% (INDELs). Carrol et al. found substantially more rejected candidates in Illumina NovaSeq 6000 data compared to Element AVITI (10). We similarly found that the proportion of rejected candidates was higher in Illumina NovaSeq X Plus across a wide range of coverages (Figure 1f).

We further evaluated the accuracy for the DeepVariant calls that passed the model evaluation. The Element AVITI was previously found to be more accurate compared to Illumina NovaSeq 6000, especially at lower genome coverage (10). For a truth set, we relied upon DeepVariant calls based on PacBio Revio data from the MM1S, OPM2 and KMS12BM cell lines (Supplementary Table 3). REH was not included, as existing PacBio data was low coverage and generated using the Sequel II instrument (12). Benchmarking analysis used hap.py (24) and was confined to only high-confidence regions on chr20 (see Methods for details). Variant calling performance was measured using the F1-score, which is the harmonic mean of recall/sensitivity and precision. Overall, our analysis showed that the variant calling performance between AVITI and NovaSeq Plus was highly comparable (Figure 1g) with AVITI only providing slightly higher F1 scores for INDELs at low coverage, particularly in the AVITI CB FS data.

Investigating these variant calls further, we looked at the performance by genomic context for SNPs and INDELs combined. For annotations on genomic context, we relied on genome-in-a-bottle (GIAB) stratifications (22). Similar to the coverage analysis, most variation relative to NovaSeq X Plus in variant calling performance was seen in shorter repetitive regions and regions with extreme GC-rates (Figure 1h). Most noticeably, the AVITI runs had increased F1 score for short (<50bp) TRs and high (75-80%) GC regions, while the F1 score in 7-11 bp GC homopolymers was reduced. The AVITI CB data also showed reduced F1 score in longer (50-149bp) 4-mer TRs compared to both AVITI CB FS and NovaSeq X Plus.

### Exploring sequencing error modes

We further investigated the sequencing error rate to evaluate the accuracy of base calls in the different samples. First, we looked at differences in the individual read compared to the reference genome. This error measure is not absolute as it also includes true variants reported as errors, but it can still highlight differences between the runs. The majority of mismatches were substitutions, making these the focus for the continued investigation (Supplementary Figure 5). Looking across cycles and reads, the NovaSeq X Plus displays a significant increase in reference-based error rate towards the end of each read (Figure 2a). A similar pattern has been observed for other Illumina instruments (42). In the AVITI run, the error rate is mostly stable across cycles, but considerably higher in read 2 for AVITI CB compared to AVITI CB FS. A similar pattern of increased error rate in read 2 could also be observed in publicly available AVITI datasets (Supplementary Figure 6a-b).

**Figure 2:**
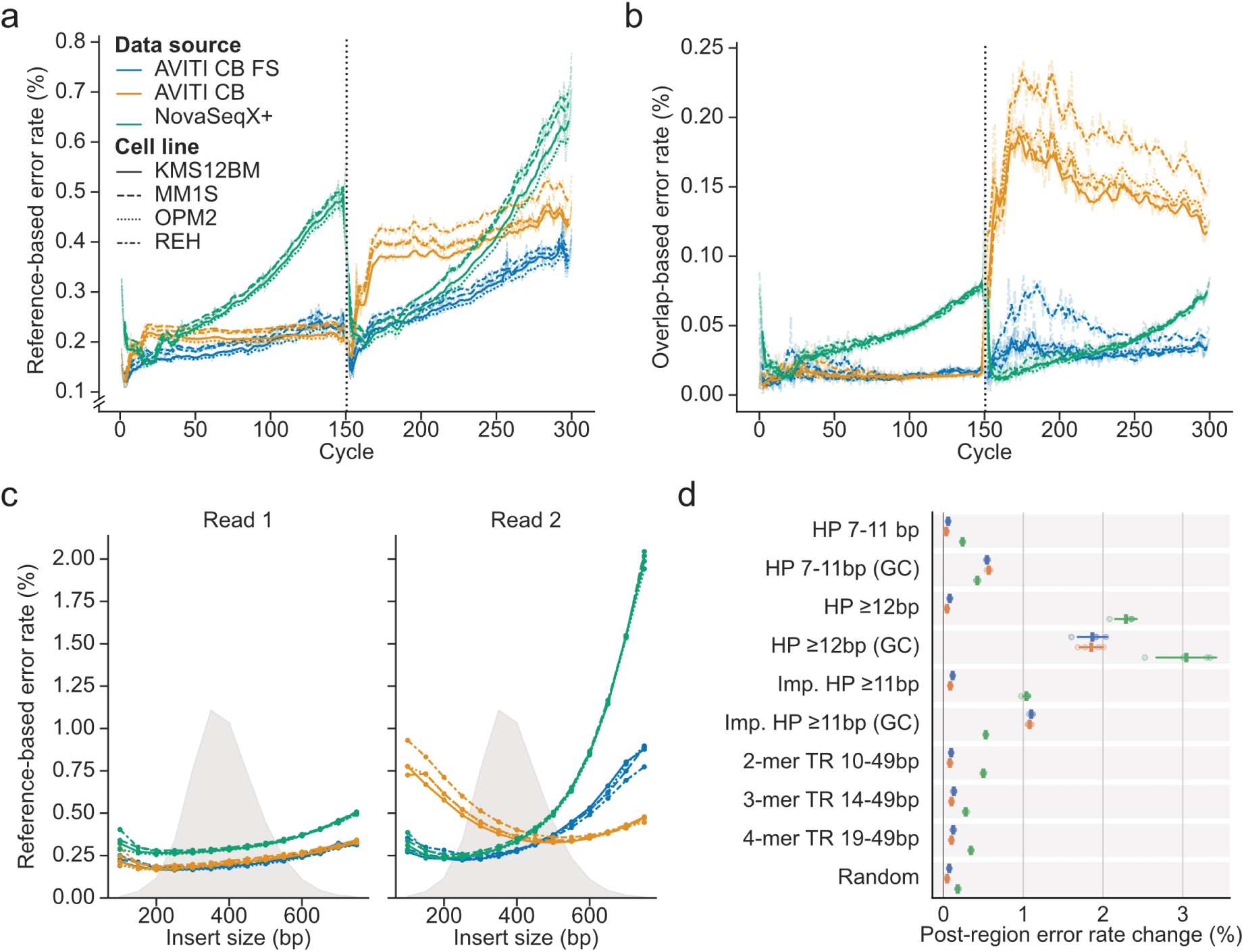
Variations in error rate in the form of substitutions across datasets. Error rate across cycles based on (**a**) difference to reference and (**b**) read overlap disagreement. The dotted line shows the boundary between reads 1 and 2. (**c**) Reference-based error rate across insert sizes. Insert sizes calculated in 50 bp bins. The gray area shows the overall reads density for each bin. (**d**) Change in error rate following small repeat regions for overlapping reads. Repeats regions acquired from GIAB stratifications (v3.5). A set of random 30 bp regions (Random) was also included for reference. Reference-based error rate computed for reads overlapping small repeats. Error rate change was calculated for the 50 bp downstream of the region relative to the 50 bp upstream. Regions composed solely of G or C bases are labelled GC. HP = homopolymer, Imp = Imperfect, TR = tandem repeat.

Instead of relying on the reference, an absolute measure of the sequencing error rate can be computed by investigating read pair overlaps (32). Read pairs with short inserts have overlapping sequences that can be used to identify errors introduced in sequencing. Importantly, these errors can be identified independently of true variants or pre-sequencing artifacts, for example caused by DNA damage. Stoler & Nekrutenko estimated the overlap-based error rate in the middle of the read for Illumina NovaSeq 6000 to a median value of 0.109% (32). For NovaSeq X Plus, this rate was estimated to be between 0.036% (OPM2) to 0.040% (REH), which is considerably lower as compared to NovaSeq 6000 rates. Using this error rate measure, the pattern of increased error towards the later cycles remains for NovaSeq X Plus, although with a lower rate overall (Figure 2b). Notably, we observe a jump in error rate for read 2 in the AVITI CB data (Figure 2b). A similar jump was also observed in one of the public AVITI CB datasets (Supplementary Figure 6C). Apart from this, we see the same trends as in the reference-based analysis, with a lower read1 error rate for AVITI.

Overlap-based error profiling is limited to pairs with inserts shorter than the combined pair read length (300 bp in this study). To investigate whether this could bias the observed jump in error rate for read2 in the AVITI CB, we profiled the reference-based error rate across insert sizes (Figure 2c). The AVITI CB sample had a clear pattern of increased error rate for short inserts in read2, which would explain the sudden increase in the overlap-based error rate (Figure 2b). A similar trend was also visible in the public Element dataset that displayed a similar jump (Supplementary Figure 6d). Investigating the AVITI CB error rate increase for short inserts in read2, we looked at the bases involved in the errors and found that the read 2 errors were primarily C>T, G>A transitions followed by C>A, G>T (Supplementary Figure 7). The same pattern could not be observed for the NovaSeqX+ and AVITI CB FS runs, where instead the error rate increased with larger insert sizes (Figure 2c). In particular, the NovaSeq X Plus samples had substantially increased error rate for ≥500 bp inserts in read2, similar to what has been observed with other Illumina instruments (43).

### Influence of repetitive regions on read error rate

Sequence context, and in particular homopolymers, are known to influence error rates in Illumina sequencing (32, 44). To investigate this further, we evaluated error rates in reads spanning repetitive regions as defined by the Genome in a Bottle (GIAB) (22). Specifically, we compared changes in error rates for bases downstream of these regions. Previous studies have shown that AVITI sequencing exhibits lower error rates downstream of long homopolymers than Illumina NextSeq 2000 and NovaSeq 6000 (5, 45). Our data show that Illumina NovaSeq X Plus has a substantially increased error rate downstream of homopolymers and tandem repeats, while AVITI maintains a relatively stable error rate in those contexts (Figure 2d). However, across GC homopolymers, AVITI shows a notable error rate increase, often exceeding that of the NovaSeq X Plus data. These elevated error rates may lead to false-positive variant calls, potentially contributing to reduced F1 scores observed in GC homopolymers of 7–11 bp in length (Figure 1h). While this highlights a potential bias in the avidity sequencing, the impact is likely limited as such GC homopolymers comprise a small fraction of the human genome (Supplementary Table 4).

### G-quadruplex motifs influence AVITI sequencing

Inspection of the datasets revealed frequent and strand-specific soft-clipping at shared locations among AVITI datasets not found in data from the other instruments (Figure 3a). Soft-clipping is a common alignment strategy used to improve alignment accuracy by omitting bases from the ends of reads that significantly diverge from the reference sequence. Notably, many of these soft-clipped loci appeared to colocalize with G-quadruplex (G4) motifs. G4 motifs can form noncanonical DNA structures composed of four guanine repeats interspersed with short loop sequences of other bases that can fold into stable secondary structures (46, 47). These motifs are common in the human genome, with one recent study finding ∼700, 000 G4 motifs (48).

**Figure 3:**
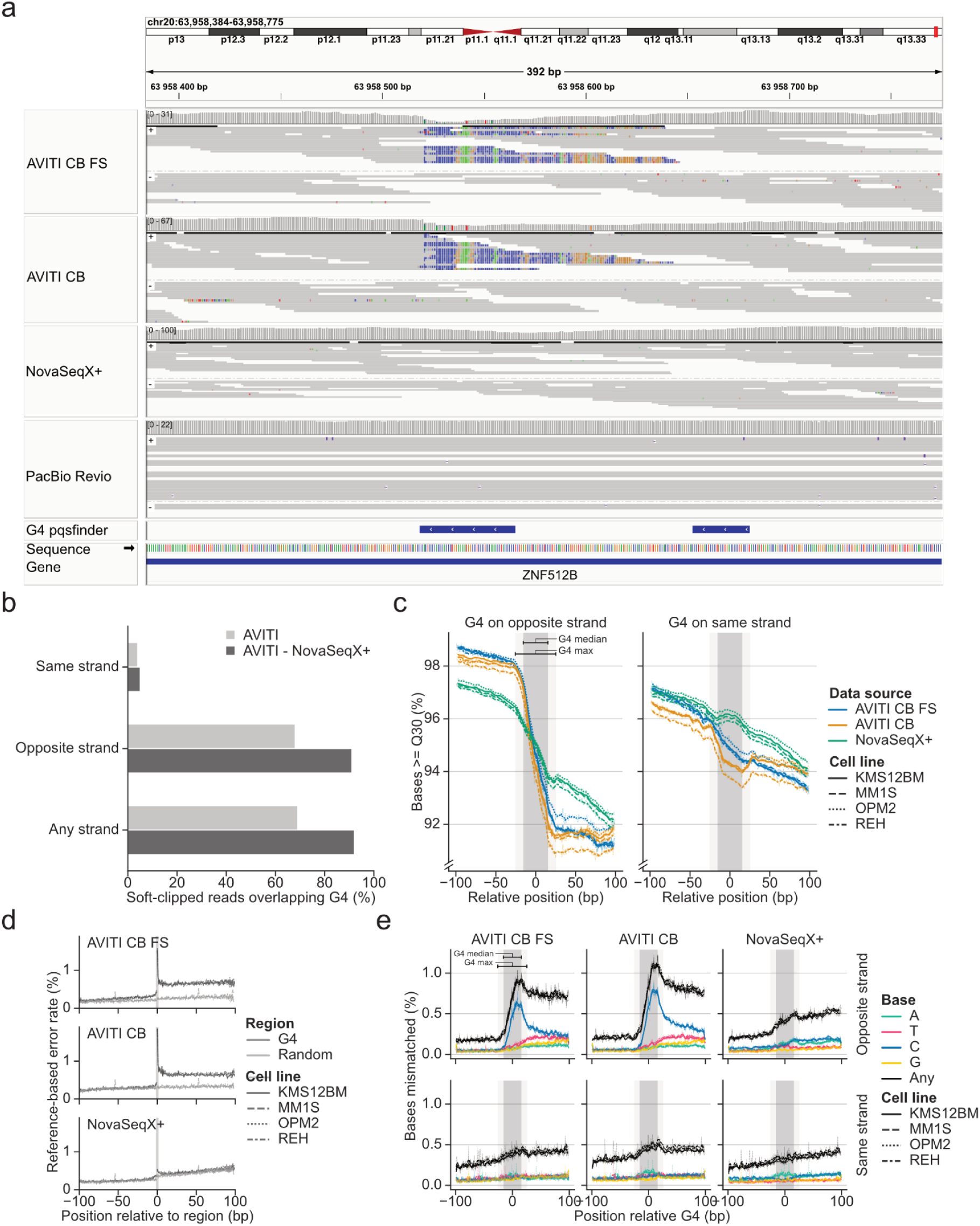
G-quadruplex (G4) related errors in Avidity sequencing. **(a)** IGV (v2.16.2) (52) example from MM1S cell line. **(b)** AVITI soft clipped reads overlap predicted G4 regions on the opposing strand. **(c)** G4 regions cause a drop in reported base quality that persists to downstream cycles. The gray region shows the median (inner) and maximum (outer) G4 span. **(d)** G4 regions cause an increase in error rate. **(e)** Erroneous base calling around G4 regions causes an abundance of C-called bases.

To explore the relationship between soft-clipping in AVITI samples and G4 motifs, we used predicted G4 regions for GRCh38 from pqsfinder (49). Reads with ≥10 soft-clipped bases were intersected by strand across AVITI datasets to identify common clipping sites. Sites overlapping across strands were excluded, and further filtering removed any commonly soft-clipped loci present in the NovaSeq X Plus data to remove shared structural variants or other artifacts across cell lines. This analysis revealed that the majority of recurrent AVITI soft-clipped sites overlapped predicted G4 motifs, and these overlaps were almost exclusively strand-specific, occurring on opposite strands (Figure 3b). Possibly this can partially explain the biases observed for AVITI in and around GC homopolymers (Figure 1e, h and 2d), as these often colocalize with G4 motifs (Supplementary Table 5).

The recurrent soft-clipping of AVITI reads overlapping G4 motifs on opposite strands suggests that avidity sequencing accuracy may be reduced in regions surrounding these noncanonical DNA structures. G4 motifs have been associated with increased error rates across multiple sequencing platforms(50, 51). To investigate this further, we analyzed base quality scores for reads overlapping predicted G4 motifs, stratified by strand. In AVITI samples, we observed a pronounced decrease in base quality following G4 motifs on the opposite strand (Figure 3c), whereas NovaSeq X Plus showed only a modest decline. Interestingly, base quality before G4 motifs was found to be elevated in AVITI reads.

This strand-specific drop in base quality in AVITI data also coincided with a rise in sequencing error rates at these loci (Figure 3d). This supports the idea that G4 structures pose a challenge for avidity sequencing. Inspection of the soft-clipped portions suggested a non-random incorporation of bases into G4 motifs (Figure 3a). Examination of mismatched bases in reads extending into G4 motifs revealed an excess of C-bases in the Element reads, a pattern not observed in Illumina data (Figure 3e). This suggests that G4 motifs may hinder accurate cycle progression in AVITI sequencing, potentially leading to erroneous variant calls (Supplementary Figure 8).

## DISCUSSION

Comparing WGS results between PCR-free genome libraries sequenced on the AVITI and NovaSeq X Plus platforms, we found several platform specific differences. The AVITI system provided lower duplication rates and also higher reported base qualities, which translated into higher mapping confidence and less spurious variant candidates. However, based on benchmarking against PacBio DeepVariant calls, overall variant calling performance was highly comparable between the AVITI and NovaSeq X Plus platforms, except for INDELs at low coverages, where the AVITI performed better. Stratifying comparisons by genomic context revealed additional differences in the platforms. Looking at both genome coverage and variant calling performance, AVITI outperformed NovaSeq X Plus in high GC regions while being inferior in GC homopolymeric regions.

Since no suitable small variant truth set exists for the cell lines employed in this study, we relied on PacBio DeepVariant calls for benchmarking. While our approach is less rigorous compared to other efforts (24, 28, 53), it should reliably detect major differences in variant calling performance between the sequencing technologies. For future studies, relying on resources such as the HG002 Q100 genome (54, 55) and the platinum pedigree (53) should help to resolve differences between the AVITI and NovaSeq X Plus in a more comprehensive manner.

Investigations of sequencing error rates showed that NovaSeq X Plus displayed increasing error rates toward read ends as expected for Illumina reads (56), while AVITI generally displayed more stable error rates. However, while AVITI error rates overall were lower in read 1, read 2 showed elevated error rates in the AVITI CB datasets. The latter was seemingly associated with shorter inserts in both our experiments and in some publicly available AVITI CB datasets. The major substitutions involved in this increased read2 error rate, C>T/G>A and C>A/G>T, could be symptomatic of DNA damage through cytosine deamination and guanine oxidation. Altogether, this points to potential issues with the read turnaround process on the AVITI instrument, although the cause is unclear.

Looking specifically at errors in repeat regions, we in line with previous results (5) found that AVITI typically maintained stable rates downstream of repetitive regions while NovaSeq X Plus showed elevated rates, particularly following longer homopolymers. The one exception to the latter was GC homopolymeric regions, where AVITI error rates sometimes exceeded NovaSeq X Plus.

Finally, AVITI sequencing was found to be particularly affected by G-quadruplex motifs that are found throughout the genome. The AVITI reads showed both pronounced quality score drops and increased sequencing errors following G4 motifs on the template strand across all samples. The sequencing errors could mainly be attributed to increased cytosine incorporation. This could be caused by polymerase stalling in the G4 guanidine repeats. Interestingly, this presents an opportunity to effectively detect G4 motifs, similar to what previously has been done using native Illumina data (57), despite lower G4 formation under normal Illumina sequencing conditions (48). G4 motifs in the template also increased quality in the bases preceding the motif (Figure 3c), which we hypothesize reflects increased polony compaction. If true, this could prove a new avenue for future quality improvements for avidity sequencing.

Overall, we conclude that AVITI and NovaSeq X Plus produced highly comparable data for human WGS. However, each platform exhibits distinct strengths and limitations, in particular relating to genomic contexts and read error modes. When selecting a sequencing platform, these differences should be carefully considered in relation to the analyzed genome or application in question.

## ACKNOWLEDGMENTS

The authors acknowledge support from the National Genomics Infrastructure (NGI) in Sweden funded by the Science for Life Laboratory, the Swedish Research Council, and the Knut and Alice Wallenberg Foundation. Additional funding was provided by the Swedish Blood Cancer Association, the Swedish Childhood Cancer Foundation, the Swedish Cancer Society and Radiumhemmets forskningsfonder. Computational resources were provided by the National Academic Infrastructure for Supercomputing in Sweden (NAISS) at UPPMAX, funded by the Swedish Research Council through grant agreement no. 2022-06725. Part of the AVITI sequencing was performed by Element Biosciences as a paid service.

## AUTHOR CONTRIBUTIONS

JN and RM conceptualized and planned the study. TM provided project administration. HF prepared PCR free Illumina libraries. ML and CN performed short-read sequencing. PL and JA performed QC and initial analysis of short-read sequencing data. JN and JH provided cells for DNA extraction. SH prepared long-read sequencing libraries and performed long-read sequencing. AA performed quality control and initial analysis on long-read sequencing data. PH integrated and curated data, performed in depth analysis, generated visualizations and wrote associated code. PH, JN and RM supervised the study. PH and RM wrote the manuscript with input from the other authors. All authors reviewed the manuscript before submission.

## CONFLICT OF INTEREST

The authors have no conflicts of interest to disclose.

## DATA AND CODE AVAILABILITY

All analysis related code including Snakemake workflows, configs, analysis scripts and notebooks are available on GitHub (<u>NationalGenomicsInfrastructure/NGI_Element_benchmark</u>) and as a persistent copy on Zenodo (https://doi.org/10.5281/zenodo.17302040). Raw Element AVITI and Illumina NovaSeq X Plus sequencing data for all cell lines were submitted to ENA under the study accession PRJEB90663. PacBio Revio data for the multiple myeloma cell lines were uploaded to ENA under the study accession PRJEB95775.

## SUPPLEMENTARY DATA

Supplementary Data are available online

## Notes

### Competing Interest Statement

The authors have declared no competing interest.

