## Supplementary Data for "Whole genome sequencing with AVITI and NovaSeq X Plus reveals comparable performance with contextual biases"

**Supplementary Table 1:** Publicly available Element datasets used. CB = Cloudbreak, FS = Freestyle.

| Dataset | Freestyle | File format | Link |
|---|---|---|---|
| Arslan et al. 2024 - HG002 (CB) | No | BAM | s3://avidity-manuscript-data/data/sentieon-bwa/20220601_PLT-03_BBS-0174-OBPA__FQD-2x150-35x/deduped.bam |
| GIAB - HG002 Long Insert (CB) | No | BAM | s3://giab/data/AshkenazimTrio/HG002_NA24385_son/Element_AVITI_20231018/HG002_GRCh38-GIABv3_Element-LngInsert_2X150_55x_20231018.bam |
| GIAB - HG002 Std Insert (CB) | No | BAM | s3://giab/data/AshkenazimTrio/HG002_NA24385_son/Element_AVITI_20231018/HG002_GRCh38-GIABv3_Element-StdInsert_2X150_81x_20231018.bam |
| Platinum Pedigree - NA12878 (CB) | No | BAM | s3://platinum-pedigree-data/data/element/mapped/GRCh38/2188-E.GRCh38.merged.sort.bam |
| GIAB - HG008-N-D Std Insert (CB) | No | BAM | s3://giab/data_somatic/HG008/Liss_lab/Element_AVITI_20240118/HG008-N-D_Element-StdInsert_61x_GRCh38-GIABv3.bam |
| Element Bio - HG002 (CB FS) | Yes | FASTQ | https://go.elementbiosciences.com/human-whole-genome-sequencing-third-party (JM-L825-HG002) |

**Supplementary Table 2:** Duplication (optical and nonoptical) rate per lane for Illumina NovaSeq X Plus libraries.

| Cell line | Lane 1 (140 pM) | Lane 2 (160 pM) |
|---|---:|---:|
| KMS12BM | 27.5% | 12.1% |
| MM1S | 29.0% | 12.7% |
| OPM2 | 28.8% | 12.8% |
| REH | 30.8% | 13.4% |

**Supplementary Table 3**: MM cell line PacBio Revio sequencing statistics.

| Cell line | Base >= Q20 (%) | N50 fragment length (bp) | Mean chr20 coverage (X) |
|---|---:|---:|---:|
| KMS12BM | 97.2% | 13,554 | 33.5 |
| MM1S | 97.3% | 13,724 | 24.4 |
| OPM2 | 97.3% | 14,569 | 30.2 |

**Supplementary Table 4:** Extent of GIAB v3.5 homopolymer stratifications in GRCh38. GC-only refers to homopolymers with repeated G- or C-bases. Bases include 5 bp on each side of each repeat (e.g. slop).

| Stratification | All bases (% of GRCh38) | GC-only bases (% of GRCh38) | % stratification GC-only |
|---|:---:|:---:|:---:|
| Homopolymers 7-11bp | 49788855 (1.6%) | 1796827 (0.1%) | 3.61% |
| Homopolymers ≥12 bp | 18344341 (0.6%) | 80541 (0.003%) | 0.44% |
| Imperfect homopolymers ≥11 bp | 44462862 (1.4%) | 2549394 (0.1%) | 5.73% |

**Supplementary Table 5:** GIAB stratifications with significant overlap with G4 motifs as computed with "bedtools fisher". Only overlaps where at least 50% of bases were shared reciprocally were considered. P-values adjusted using Bonferroni method and only stratifications values below 0.01 kept. HP = homopolymer, Imp. HP = imperfect HP.

| Stratification | Regions | Overlapping G4 | % overlapping | Odds ratio |
|---|---:|---:|---:|---:|
| HP ≥21bp (GC) | 66 | 64 | 97.0% | 1,009.8 |
| Imp. HP ≥21bp (GC) | 3,439 | 2,993 | 87.0% | 213.4 |
| HP ≥12bp (GC) | 3,398 | 2,673 | 78.7% | 145.0 |
| Imp. HP ≥11bp (GC) | 107,713 | 77,378 | 71.8% | 106.2 |
| HP 7-11bp (GC) | 101,792 | 30,204 | 29.7% | 19.0 |
| HP 4-6bp (GC) | 12,048,931 | 613,434 | 5.1% | 3.7 |
| Imp. HP ≥11bp | 1,678,630 | 76,546 | 4.6% | 1.8 |
| HP ≥7bp + Imp. HP ≥11bp | 3,819,656 | 95,925 | 2.5% | 1.0 |

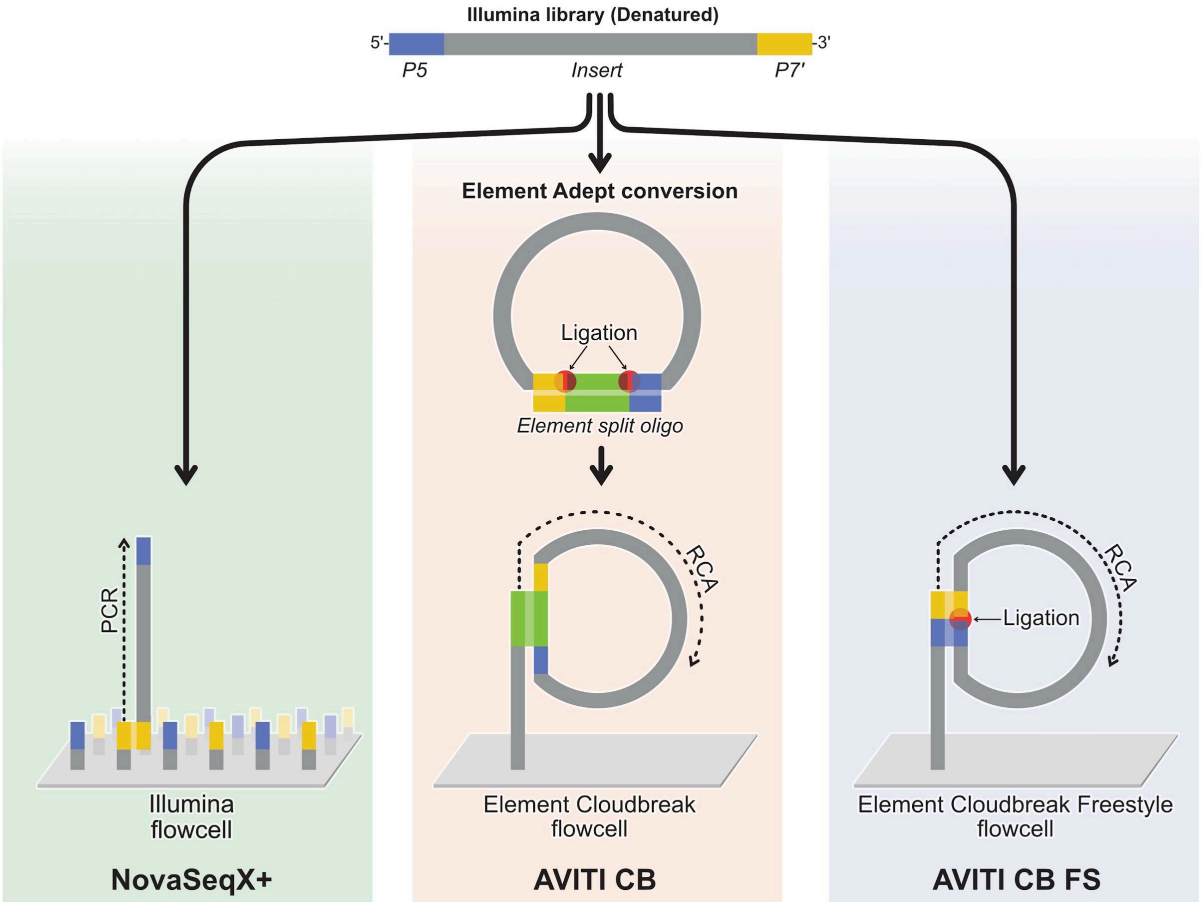


**Supplementary Figure 1:** Schematic illustration of the different Illumina NovaSeq X Plus and Element AVITI sequencing workflows utilized in the study. CB = Cloudbreak, FS = Freestyle.

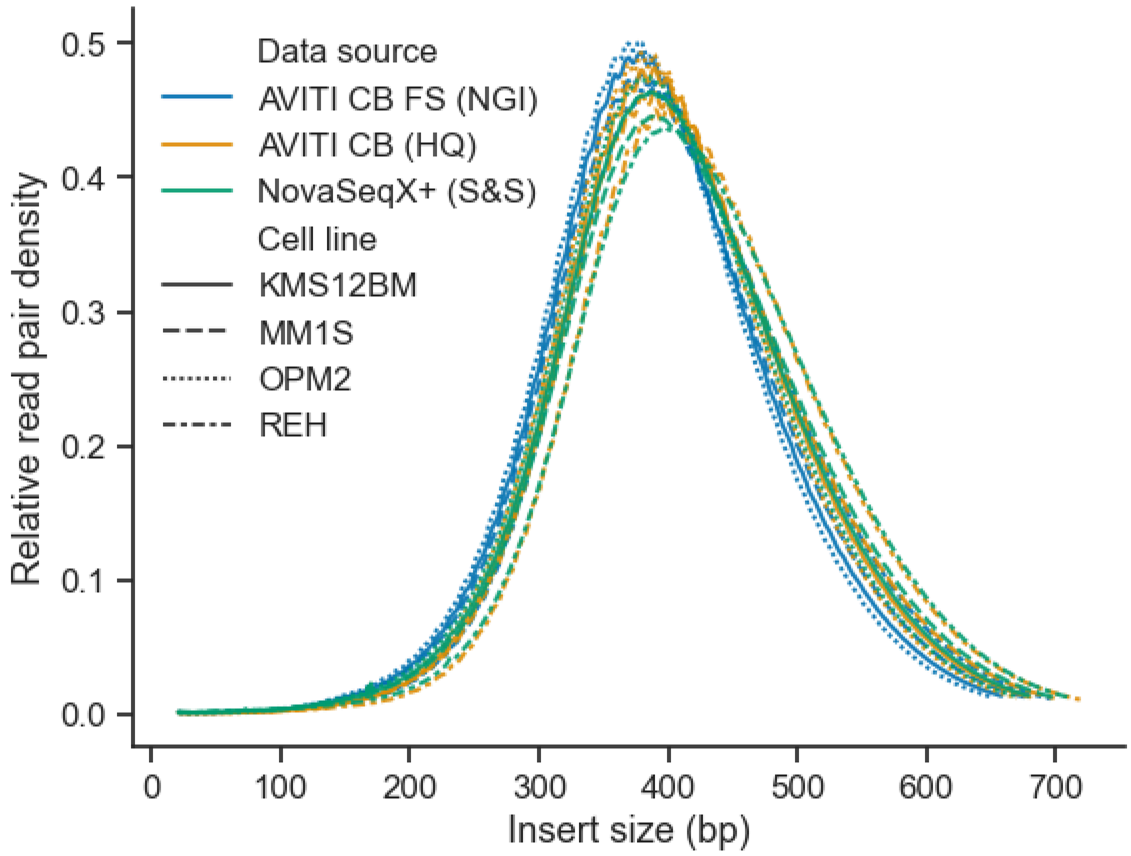


**Supplementary Figure 2:** Insert size distribution for the mapped libraries.

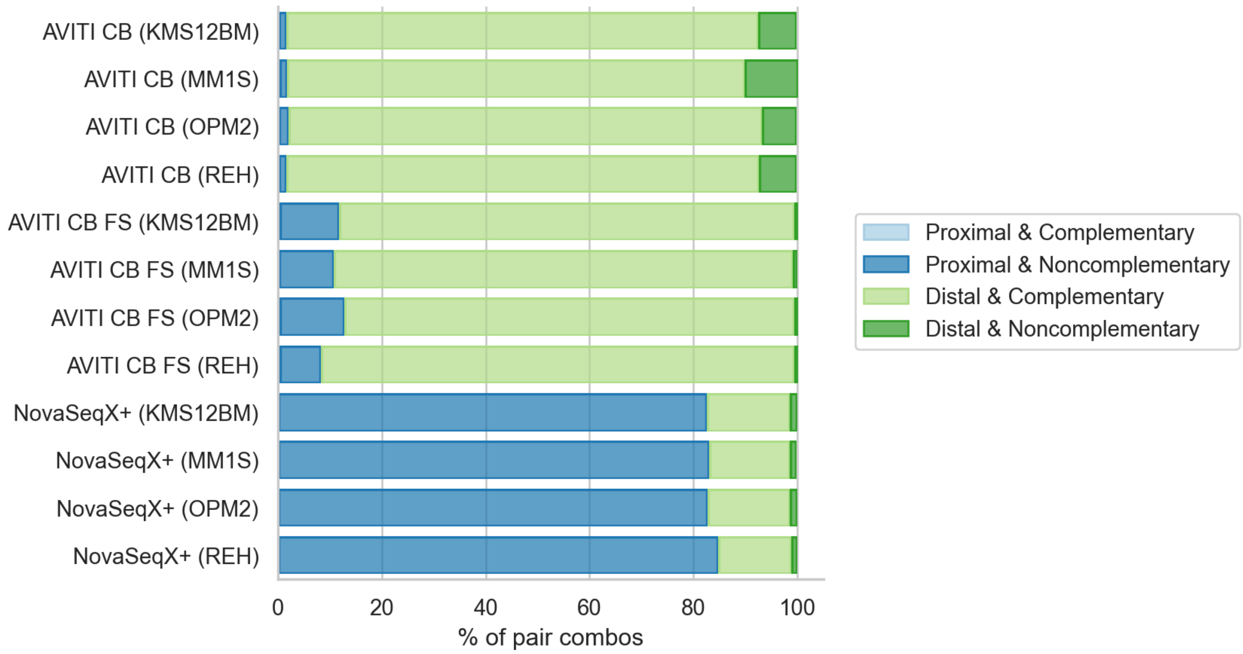


**Supplementary Figure 3:** Duplicate read pair classifications based on physical proximity on flow cell and complementarity. Read pairs mapped to chr20 sharing mapping positions and pair orientation were grouped, and within each group the pair combinations were classified. Pair combinations were classified as “Proximal” if their physical distance was less than 100 distance units (i.e. are optical duplicates). Pair combinations were further classified as “Complementary” if the pairs were in opposite directions with regard to read1 and 2 (i.e. representing the complementary strands of the same molecule).

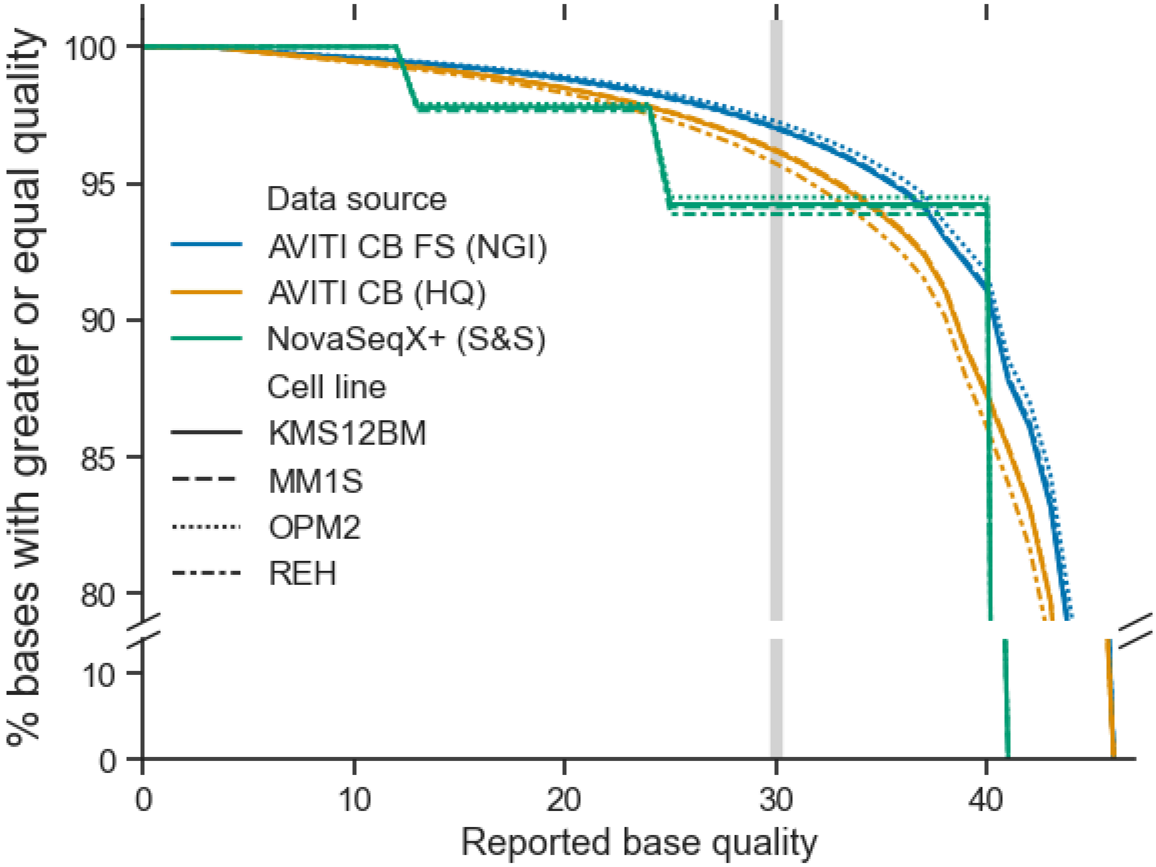


**Supplementary Figure 4**: Reported base quality distribution shown as the percentage of bases with a certain Q-score or greater. Illumina NovaSeq X Plus output Q-scores in four bins producing the stepped graph shown here. The highlighted area in grey indicates Q30.

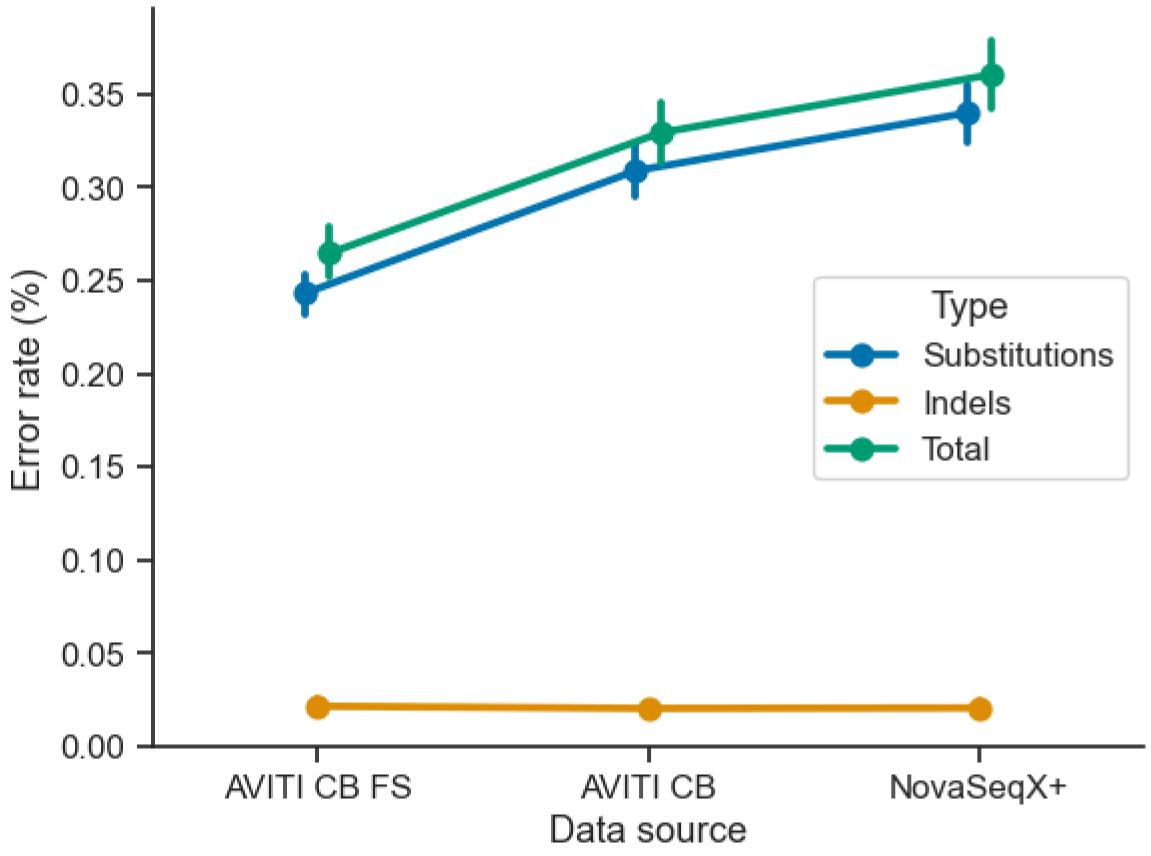


**Supplementary Figure 5**: Error rate as the percentage of bases mismatched to the reference is primarily driven by substitutions. Note that the mismatches include true variants.

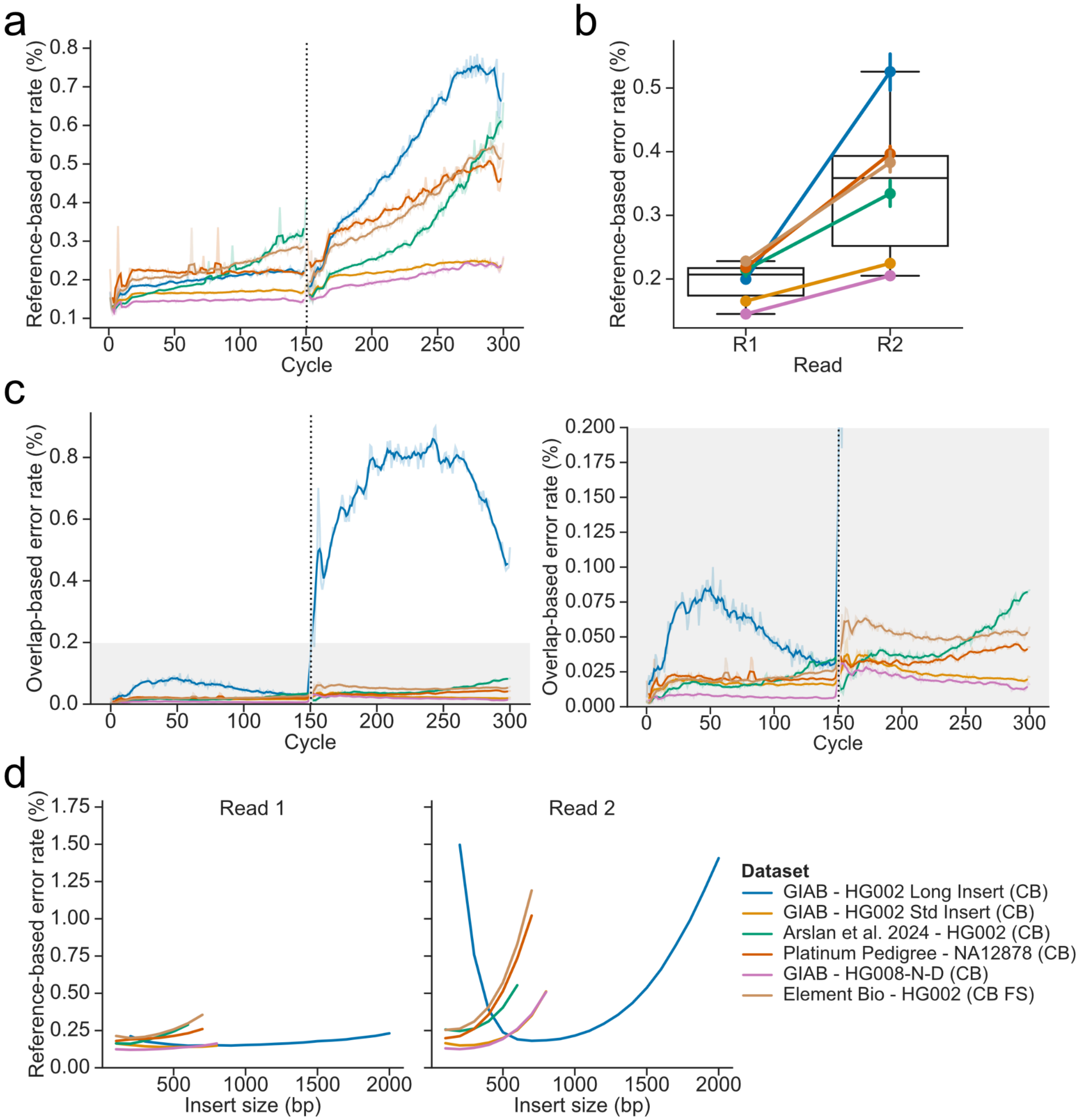


**Supplementary Figure 6**: Error rates in public Element AVITI datasets. (**a**) Error rate across cycles based on mismatches to the reference. The dotted line shows the boundary between reads 1 and 2. (**b**) Change in average error rate between read 1 and 2. (**c**) Error rate across cycles based read pair overlap mismatches. The right plot is a zoomed in version of the left plot within the area highlighted in grey. (**d**) Error rate across reads and insert sizes. Insert sizes calculated in 50 bp bins.

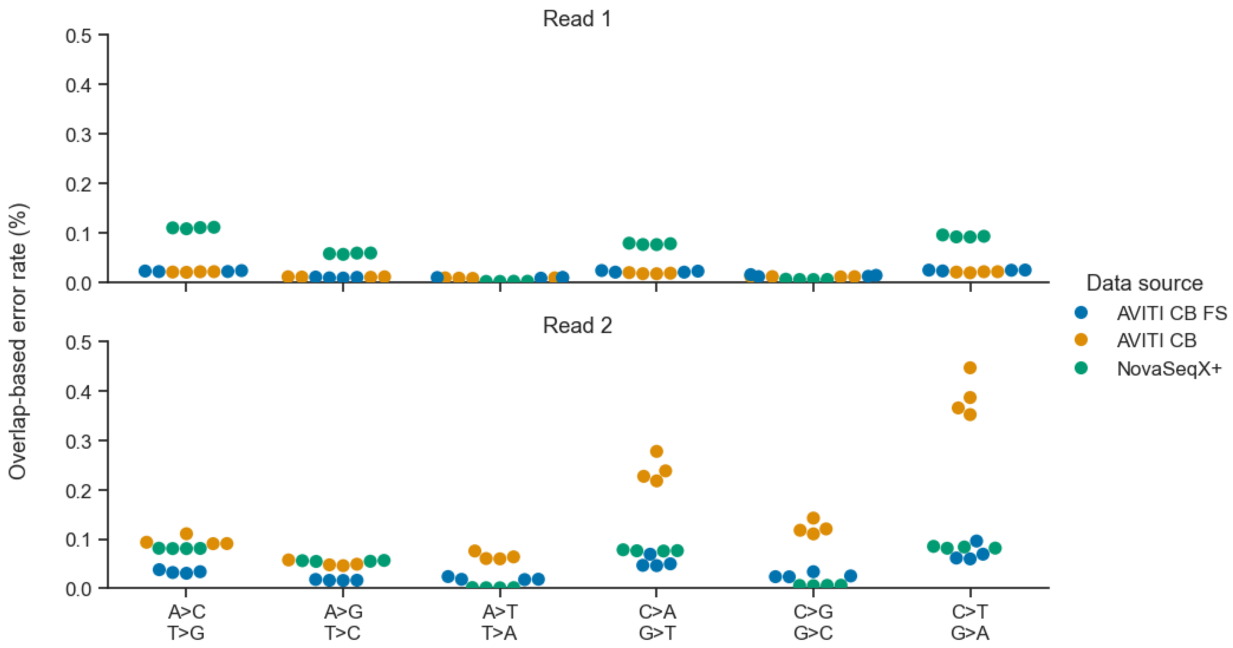


**Supplementary Figure 7**: Error rate by sequence context and read. Error rate calculated using fraguracy based on overlapping bases in read pairs. Note that only short fragments with an insert size <300 are included here.

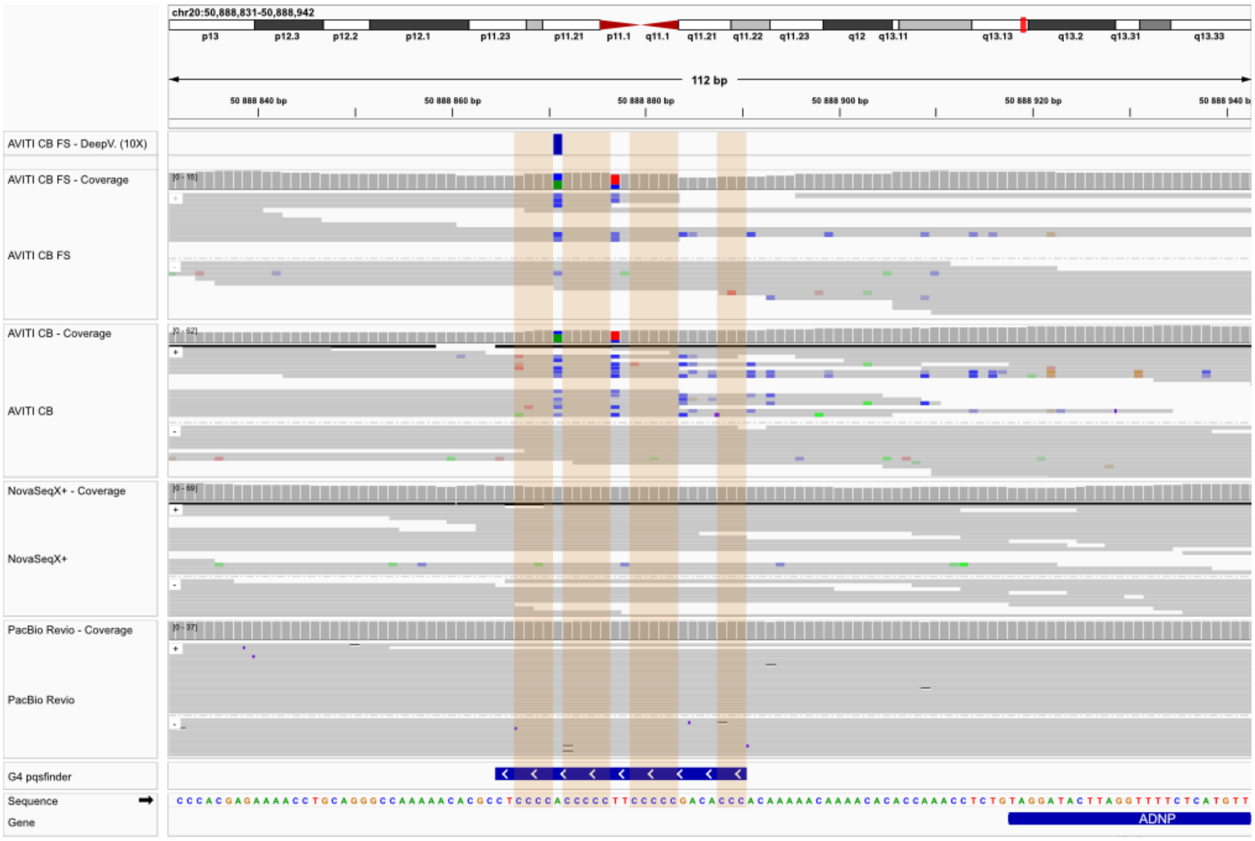


**Supplementary Figure 8:** Example of cysteine base-call errors in AVITI sequencing data related to G4 motif on opposing strand. Highlighted sections in orange show polyG stretches of G4 motif. Top row shows DeepVariant calls with AVITI CB FS at 10X coverage, generating a false positive substitution A>C in the G4-motif loop. Image shows data from MM1S cell line and was generated using IGV (v2.16.2) with soft-clipped bases hidden.
